# Rapid shallow-water saturation and deep-water expansion of an invasive freshwater ecosystem engineer in a deep European lake

**DOI:** 10.64898/2026.06.26.734794

**Authors:** Linus Hofstetter, Thomas M. Müller, Mathys Bourqui, Lyubov E. Burlakova, Zoe C. Cristante, Alexander Y. Karatayev, Sophie Kessler, Anita Narwani, Joana L. Santos, Lars Sturm, Noemi Wellauer, Piet Spaak, Alexandra A.-T. Weber

## Abstract

Quagga mussels (*Dreissena rostriformis bugensis*) are ecosystem engineers that can alter nutrient cycling, benthic-pelagic coupling, and food-web structure in deep lakes. Although their invasion trajectories are well documented in the Laurentian Great Lakes in North America, depth-specific population dynamics remain poorly resolved in recently invaded European perialpine lakes. We analyzed five annual lake-wide surveys (2021-2025) from 54 stations spanning 2.4-253 m depth in Lake Constance to quantify changes in quagga mussel density, biomass, and shell-length distribution. Contrary to expectations of lake-wide exponential growth, shallow-water populations (< 20 m) showed no significant increase during the study period and appear to have reached carrying capacity before monitoring began. In contrast, densities increased monotonically at intermediate depths (40-125 m), indicating ongoing expansion into deeper strata. Mean shell length declined with depth, and size distributions in shallow waters shifted toward larger individuals, consistent with a transition from active recruitment to somatic growth of established mussels. Compared with the Laurentian Great Lakes, Lake Constance already has substantially higher shallow-water biomass, whereas deeper invasion trajectories are broadly similar. These results show that quagga mussel invasion in deep European lakes can combine rapid littoral saturation with slower profundal expansion, complicating direct transfer of predictions from the Great Lakes. Continued depth-stratified monitoring will be essential for anticipating future ecosystem effects in perialpine lakes.

## Introduction

Biological invasions are major drivers of ecological degradation in aquatic and terrestrial ecosystems, often causing biodiversity loss and altered ecosystem function (Pyšek et al. 2020; Gallardo et al. 2024). Their economic costs have risen sharply with increasing global connectivity (Cuthbert et al. 2021; Diagne et al. 2021). In aquatic systems, invasive filter feeders are especially consequential because they can exert strong top-down control on phytoplankton, nutrient dynamics, and food-web structure (Higgins and Vander Zanden 2010; Mayer et al. 2014; Sánchez et al. 2016). Zebra mussels (*Dreissena polymorpha*) and quagga mussels (*Dreissena rostriformis bugensis*) are among the most impactful freshwater invaders in the Northern Hemisphere. Through filtration and biodeposition, *Dreissena* spp. can increase water clarity, redistribute primary production, alter phosphorus cycling, and damage water-use infrastructure through biofouling (Connelly et al. 2007; Karatayev et al. 2015; Li et al. 2021).

Population development of dreissenids has been studied for more than 30 yr in Lakes Erie, Huron, Michigan, and Ontario, hereafter the Laurentian Great Lakes. Zebra and quagga mussels were introduced to the Great Lakes in the late 1980s, rapidly colonized shallow waters, and subsequently diverged in their habitat use and population trajectories (Karatayev and Burlakova 2025b). Zebra mussels are often dominant in shallow, hard-substrate habitats during early invasion, whereas quagga mussels can settle on both hard and soft sediments and expand into deeper waters (Karatayev and Burlakova 2025c). In depths > 40 m, quagga mussels may form the profunda morphotype, characterized by flatter shells, reduced pigmentation, and more rounded ventral edges (Dermott and Munawar 1993; Pavlova 2012). This ability to occupy profundal habitats allows quagga mussels to colonize entire lake beds, dominate benthic biomass, and shift biomass maxima toward deeper strata over time (Burlakova et al. 2018; Karatayev and Burlakova 2025b).

Quagga mussels are now spreading rapidly across Europe. After expansion in Russia in the 1990s, they were recorded in Western Europe in the Netherlands in 2006 and in Belgium in 2009 (Molloy et al. 2007; Matthews et al. 2013). Since then, they have been detected in major European rivers and large lakes, including Lake Geneva, Lake Bourget, Lake Garda, and recently several Swiss perialpine lakes (De Ventura et al. 2017; Kraemer et al. 2023). In Lake Constance, the first quagga mussels were reported in 2016; within one year, the species had been detected across the lake (Haltiner et al. 2022). As in the Great Lakes, quagga mussels now occur down to the deepest parts of Lake Constance, while zebra mussels are largely restricted to a few shallow nearshore sites.

A recent cross-lake synthesis shows that *Dreissena* impacts do not scale with depth alone. It was shown that lake area strongly affected the timing and magnitude of ecosystem responses, with impacts appearing more slowly but persisting longer in the Laurentian Great Lakes than in smaller inland lakes (Karatayev et al. unpubl.). The same synthesis also identified Lakes Constance and Geneva as important deviations from typical dreissenid responses, because strong mussel populations have not yet produced clear increases in transparency or declines in phytoplankton. These findings raise an unresolved question for recently invaded perialpine lakes: whether their population trajectories will follow the deep-water expansion observed in the Laurentian Great Lakes, the faster dynamics of smaller inland lakes, or a distinct path shaped by morphometry, productivity, and mixing regime.

Lake Constance, located at the border of Switzerland, Austria, and Germany, is one of the largest and deepest lakes in Europe (Petri 2006; Rinke et al. 2009). The lake has undergone strong re-oligotrophication since the late 20th century, and climate warming has reduced the frequency of full water-column mixing events (Fink et al. 2015; Murphy et al. 2018). Because deep perialpine lakes share key features with the Laurentian Great Lakes, Great Lakes data, together with early data from Lake Constance and other deep European lakes were used to project a 9- to 20-fold increase in quagga mussel biomass in deep perialpine lakes over the coming three decades (Kraemer et al. 2023). We therefore expected quagga mussel density and biomass to increase across the lake, with the strongest growth at intermediate depths. We asked: (1) how are quagga mussel density and biomass changing during early invasion in a deep perialpine lake; (2) how do population dynamics vary with depth; and (3) how do trajectories in Lake Constance compare with those in the Laurentian Great Lakes? To address these questions, we analyzed five annual lake-wide surveys across 54 stations spanning the full depth gradient of Lake Constance.

## Materials and methods

### Data collection

Samples were collected annually in autumn from 2021 to 2025. Sediment was sampled at 28 stations in 2021 and at 54 stations from 2022 to 2025 (Fig. 1A; Supporting Information Table S1). Stations ranged from 2.4 m depth to the deepest point of the lake at 253 m and were arranged along four transverse transects, one longitudinal transect through the deepest parts of the lake, and additional stations in Lower Lake Constance and the Bay of Bregenz to represent major lake regions (Fig. 1A; Supporting Information Fig. S1).

**Figure 1.**
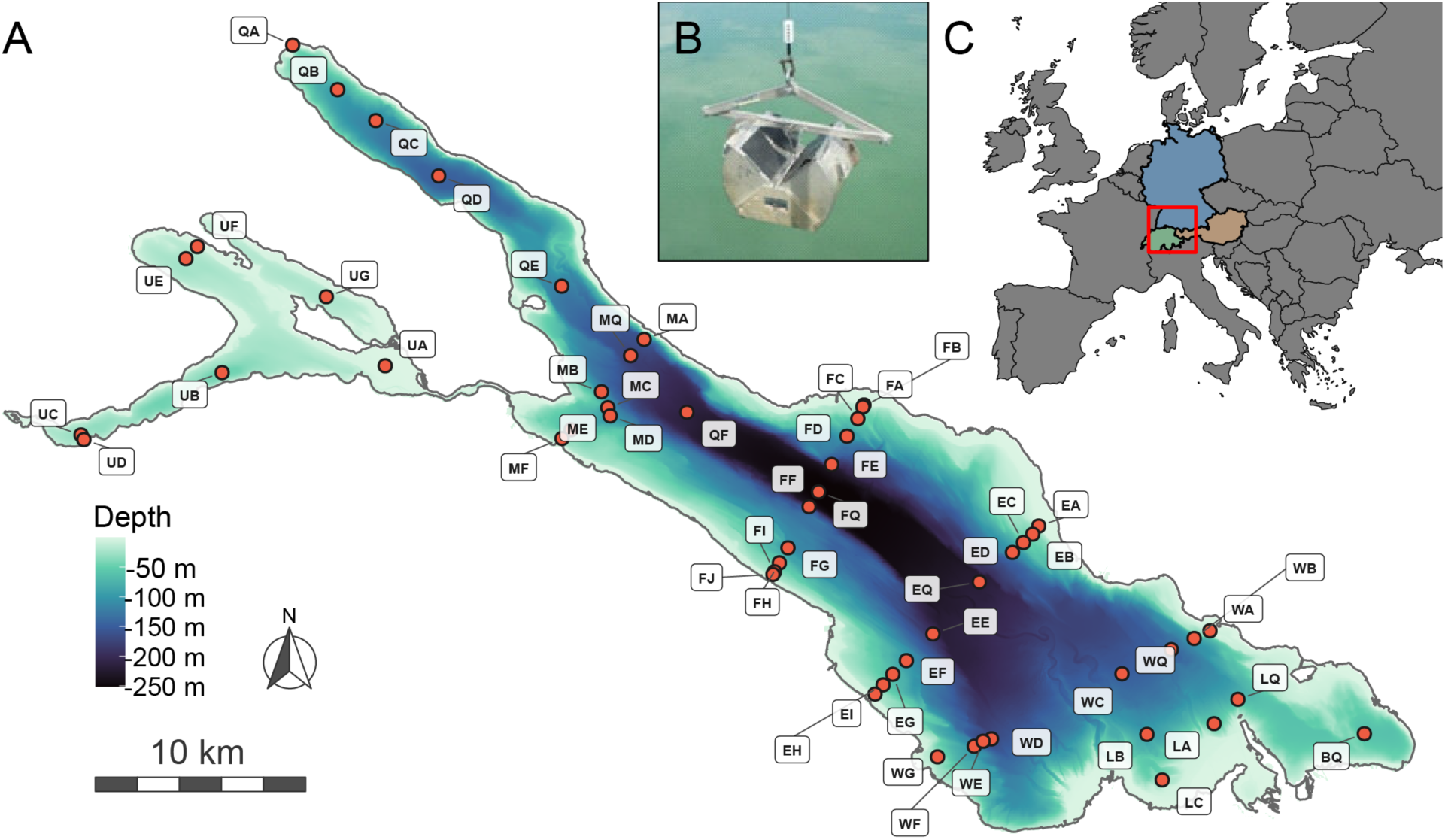
Sampling design in Lake Constance. (A) Location of the 54 sampling stations, shown as red points and labelled with two-letter station identifiers; colors show lake bathymetry. (B) Ponar sediment grab used for replicate sediment sampling at each station. (C) Location of Lake Constance in central Europe, bordering Austria (orange), Germany (blue), and Switzerland (green).

At each station, three replicate samples were collected with a Ponar sediment grab (area = 0.052 m²; Fig. 1B) (Mudroch and MacKnight 1994). Sediment was washed through a 1-mm mesh sieve and stored at -18°C until processing. In the laboratory, quagga mussels were counted to estimate density, and shell length was measured to the nearest 0.01 mm with a digital caliper (Garant HCT 412780). For large samples, randomly selected subsamples were measured until at least 150 mussels had been measured. Zebra mussels were identified and measured separately. Because quagga mussels overwhelmingly dominated the dreissenid assemblage in Lake Constance, mussel bodies without shells were counted as quagga mussels. Mussels < 5 mm shell length were counted in all years but were measured inconsistently in 2021 and 2022.

In addition to Ponar sampling, mussel densities in 2021 and 2022 were estimated from images collected with a Benthic Imaging System (BIS) (Karatayev et al. 2021a). The BIS consists of a metal frame equipped with GoPro cameras and dive lamps and was lowered to the lake bed three times at each station. Images covering 50 × 50 cm were extracted from the video footage, and individual mussels were counted. Because only 28 stations were sampled with Ponar grabs in 2021, density estimates for the remaining stations were derived from BIS image counts collected during the same sampling campaign. A detailed description of the field protocol is provided by Flämig et al. (2024).

### Biomass estimation and analysis

Ash-free dry weight (AFDW) was estimated from shell length using measurements made in 2022 on 761 quagga mussels from Lake Constance. Mussels spanning the observed size range were measured, dried for 48 h, combusted at 550°C for 3 h, and weighed. A regression between shell length and AFDW was fitted, and depth-specific coefficients were estimated for four depth categories (Supporting Information Fig. S2, Table S2, Formula S1) following Seitz (2024). For the estimate of total wet mass reported in the Results, AFDW was converted to wet weight using a factor of 37, that was experimentally determined at Buffalo State University (Burlakova, pers. comm.).

Because BIS and Ponar estimates were comparable for mussels > 5 mm shell length in previous Great Lakes work (Karatayev and Burlakova 2025a), we evaluated their relationship for stations sampled with both methods in Lake Constance. For 2021 and 2022 stations with both BIS and Ponar data, a regression between BIS counts and Ponar densities for mussels > 5 mm shell length was fitted (n = 80, slope = 0.927, R² = 0.696, p < 0.001; Supporting Information Fig. S3). This regression was used to predict 2021 densities at the 26 stations without Ponar samples. Because mussels < 5 mm were not measured consistently in 2021 and 2022, all density and biomass analyses were restricted to mussels > 5 mm shell length.

To examine depth-specific dynamics, stations were grouped into five depth strata: < 20 m, 20-40 m, 40-75 m, 75-125 m, and > 125 m. Each stratum contained eight to 13 stations (Supporting Information Fig. S1). Because the time series included only five annual observations and the data were right-skewed and zero-inflated, we primarily used non-parametric approaches. Trends in density and biomass within each depth stratum were assessed using Mann-Kendall trend tests. Whole-lake means were estimated by weighting depth-stratum means by the proportion of three-dimensional lakebed area in each stratum. Bathymetric data were obtained from PANGAEA for Lake Constance (Felden et al. 2023) and from the U.S. National Geophysical Data Center for Lakes Huron, Michigan, and Ontario (National Geophysical Data Center 1996, 1999a, 1999b). Depth-dependent changes in shell-length distributions were evaluated with permutation-based omnibus tests comparing 2023, 2024, and 2025 within each depth stratum. The years 2021 and 2022 were excluded from this analysis because mussels < 5 mm shell length were not measured consistently. Observations were randomly permuted 9999 times among year groups. Where omnibus tests were significant, pairwise post hoc permutation tests compared skewness among years within each depth category, with p-values Bonferroni-corrected for three comparisons. Skewness was calculated using the moments package in R (Komsta and Novomestky 2022). Station-level biomass change was assessed with simple linear models fitted to log-transformed biomass over the five sampling years. Biomass in Lake Constance was compared with data from Lakes Huron, Michigan, and Ontario, which are the Great Lakes most comparable in depth and stratification. Lake Erie was excluded because it is much shallower, and Lake Superior was excluded because it remains largely unaffected by quagga mussels. All analyses were performed in R version 4.5.2 (R Core Team 2025); packages used are listed in Supporting Information Table S3. Generative artificial intelligence (Claude Sonnet 4.6 (Anthropic) and ChatGPT - 5.5 (OpenAI)) were used to assist with writing and tidying of R scripts, and for outlining and grammar correction during manuscript drafting.

## Results

### Shell length declines with depth

Quagga mussels in Lake Constance reached maximum shell lengths of approximately 25 mm, and mean shell length declined with increasing depth (Fig. 2A-F). In 2025, mean shell length was 11.78 mm at < 20 m, 9.98 mm at 20-40 m, 7.72 mm at 40-75 m, 6.00 mm at 75-125 m, and 5.11 mm at > 125 m (Fig. 2F). Omnibus permutation tests showed that the right skew of shell-length distributions decreased significantly between 2023 and 2025 in four of the five depth strata (p < 0.01), with the exception of the 20-40 m stratum (p = 0.281). The largest relative reduction occurred in the shallowest stratum, where skewness decreased by 73%, from 0.61 in 2023 to 0.16 in 2025 (p < 0.001; Supporting Information Fig. S4). Reductions were more moderate at greater depths: skewness decreased by 22% at 40-75 m (1.42 to 1.11; p < 0.001), 20% at 75-125 m (1.74 to 1.39; p < 0.001), and 28% at > 125 m (2.08 to 1.49; p = 0.007).

**Figure 2.**
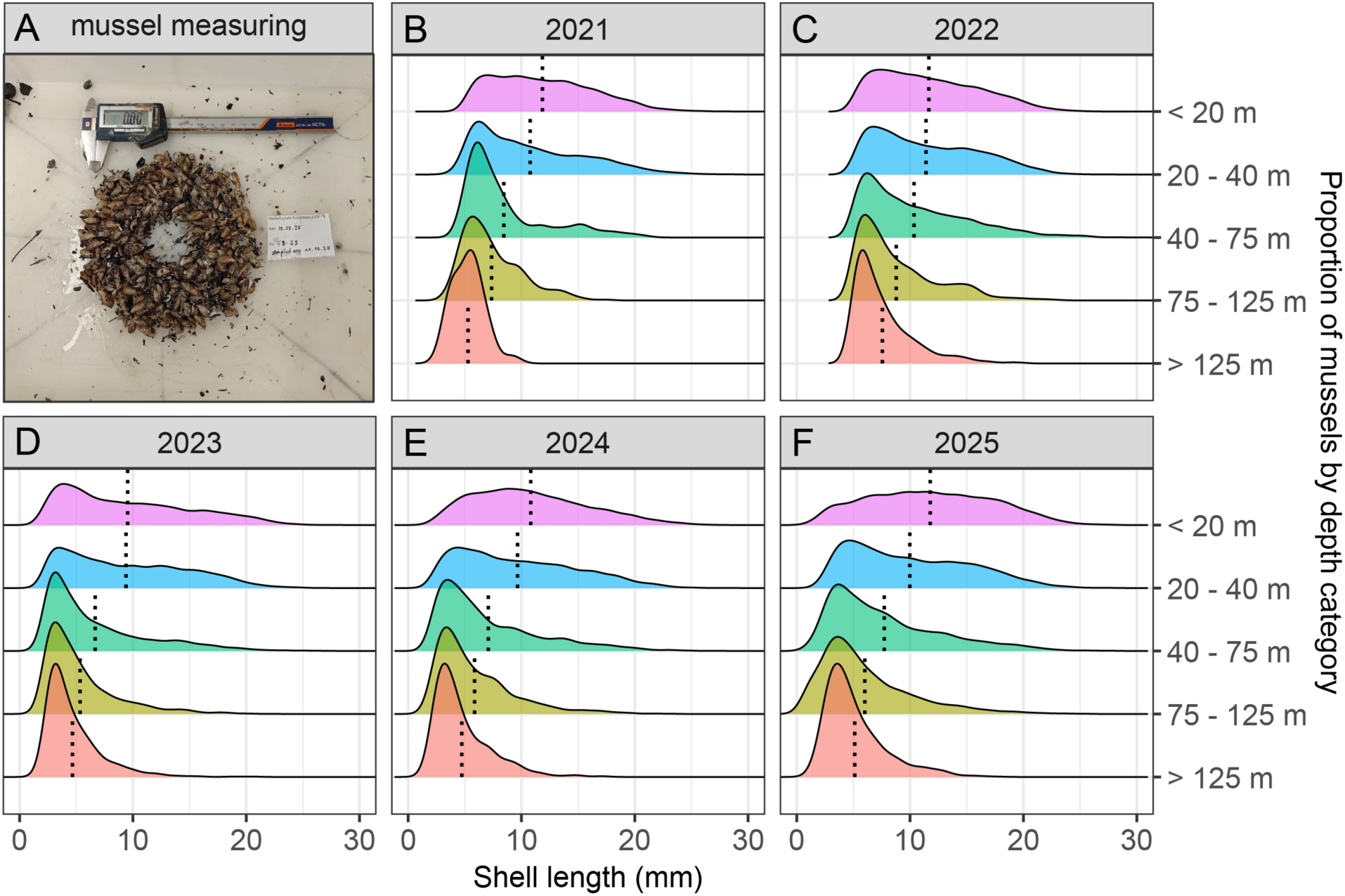
Shell-length distributions of quagga mussels by depth stratum and year. (A) Example of mussel measurement in the laboratory. (B-F) Proportion of mussels within shell-length classes in 2021-2025 for five depth strata. Dotted vertical lines indicate mean shell length within each stratum. Mussels < 5 mm shell length were not measured consistently in 2021 and were not measured in 2022.

### Density and biomass increase at depth

Both mussel density and biomass decreased with increasing depth (Fig. 3). Shallow areas < 20 m appeared to stabilize below approximately 10,000 individuals m⁻² and 35 g AFDW m⁻² for mussels > 5 mm shell length, although uncertainty was high. Mann-Kendall tests showed no significant trend in biomass or density in the shallowest stratum over the study period (biomass: p = 0.462, τ = -0.4; density: p = 0.221, τ = -0.6). Trends in the 20-40 m stratum were also not significant (biomass: p = 0.086, τ = 0.8; density: p = 0.462, τ = 0.4).

**Figure 3.**
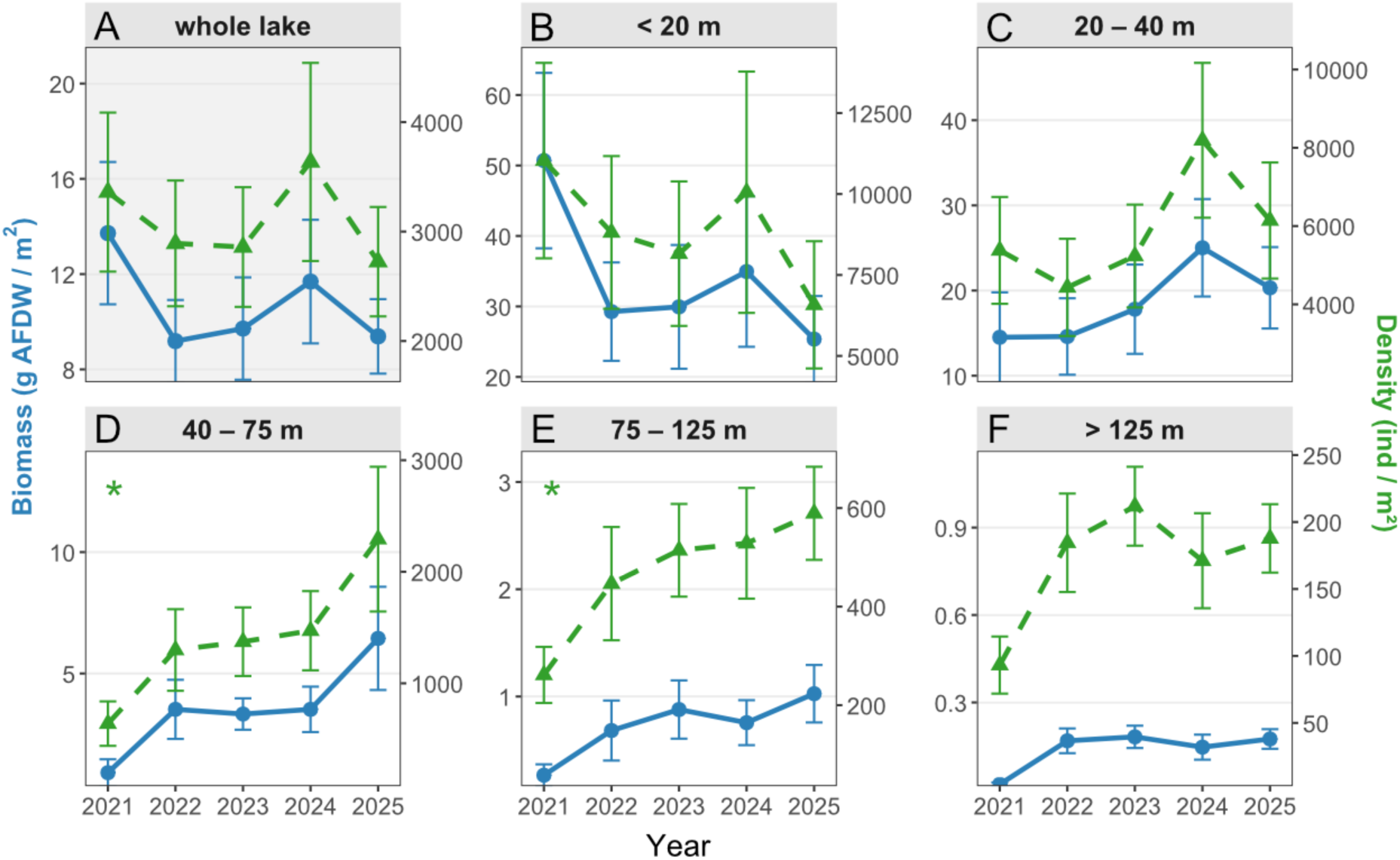
Mean biomass and density of quagga mussels > 5 mm shell length from 2021 to 2025. Biomass is shown as ash-free dry weight (AFDW, blue) and density as individuals m⁻² (green). (A) Lake-wide means weighted by the proportion of three-dimensional lakebed area within each depth stratum. (B-F) Mean values by depth stratum. Error bars show standard errors. Stars indicate significant Mann-Kendall trends.

Significant population increases occurred at intermediate depths. Density increased monotonically across all sampling years in both the 40-75 m and 75-125 m strata (p = 0.027, τ = 1.0 for both). Biomass also showed positive trends, but these were not significant at the 0.05 level (40-75 m: p = 0.221, τ = 0.6; 75-125 m: p = 0.086, τ = 0.8). In the deepest stratum (> 125 m), neither density nor biomass changed significantly (p = 0.462, τ = 0.4 for both). Weighted across the whole lake, no significant change in total density or biomass was detected over the five-year period. At a mean biomass of approximately 10 g AFDW m⁻² and a lake area of 535 km², we estimate total quagga mussel AFDW at approximately 5250 tonnes in 2025. Using a conversion factor of 37, this corresponds to approximately 190,000 tonnes wet weight. During the 2025 campaign, zebra mussels were found at only six of the 54 sampled stations, and at four of these stations only one zebra mussel individual was sampled.

### Station-level biomass trends

Linear models fitted to station-level biomass showed no significant change at 43 of 54 stations (Fig. 4). Of the 11 stations with significant trends, nine had positive slopes and two had negative slopes. Only two stations, UG and UF in oxygen-deprived parts of Lower Lake Constance (Güde et al. 2009), remained quagga-free throughout the study period. No station < 20 m depth showed a significant biomass increase, and one shallow station (WG) showed a weak but significant decrease. Although many station-level trends were not significant, likely because the time series contained only five years, 20 of 26 stations deeper than 75 m showed positive slopes (Supporting Information Fig. S5).

**Figure 4.**
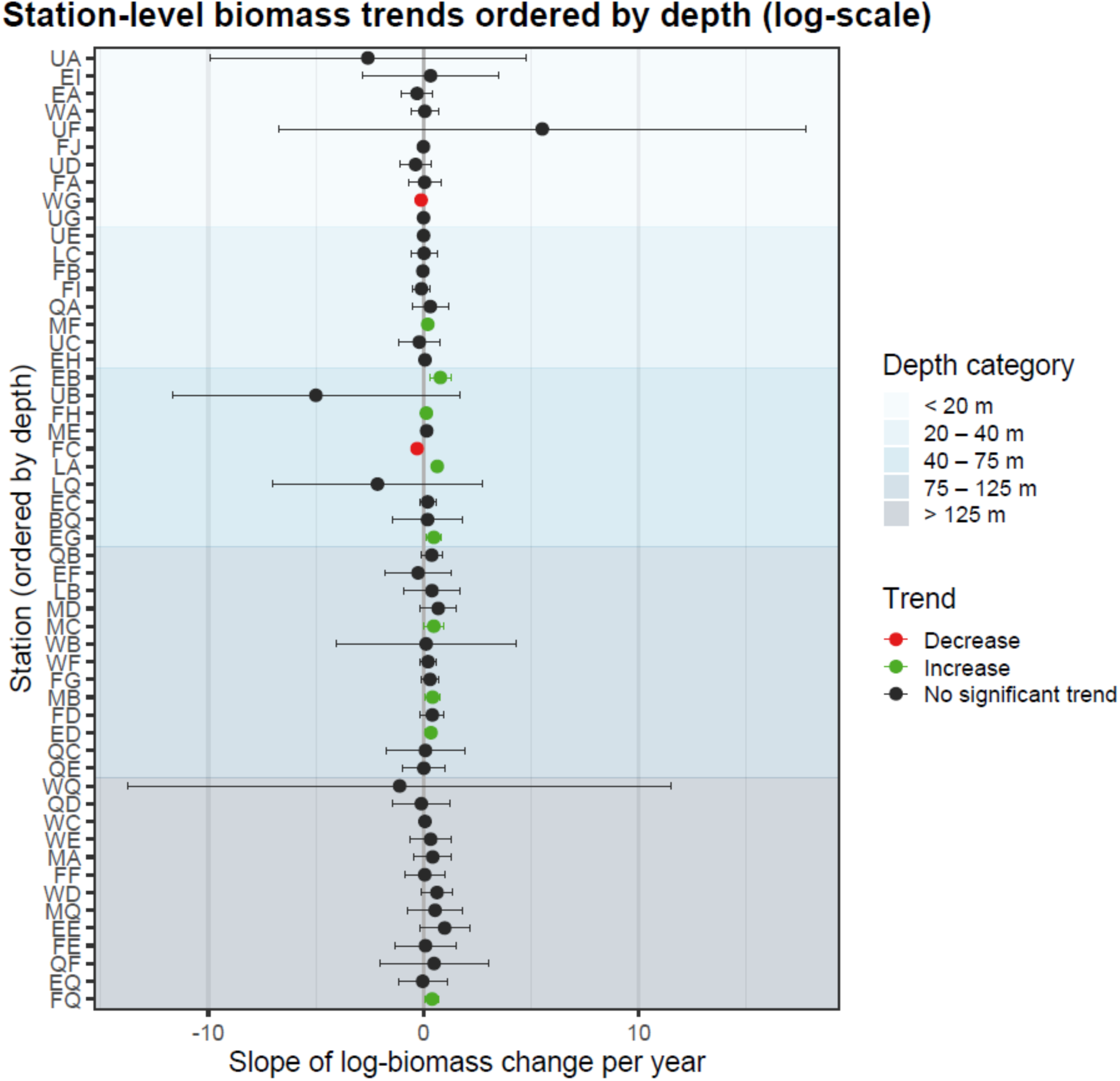
Station-level biomass trends from 2021 to 2025. Points show slopes from linear models fitted to log-transformed biomass at each station, ordered by depth. Black points indicate non-significant trends, green points significant positive trends, and red points significant negative trends. Horizontal bars show uncertainty around slope estimates, and background shading indicates depth strata.

### Comparison with the Laurentian Great Lakes

Weighted whole-lake biomass in Lake Constance was higher than biomass in Lakes Huron, Michigan, and Ontario at comparable times after first detection of quagga mussels (Fig. 5A). This difference was driven by high shallow-water (< 20 m) biomass in Lake Constance: Mean biomass for mussels > 5 mm shell length reached its maximum at 2.79 g AFDW m⁻² in Lake Huron, 4.54 g AFDW m⁻² in Lake Michigan, and 13.53 g AFDW m⁻² in Lake Ontario, compared with > 40 g AFDW m⁻² in Lake Constance in 2021. At depths > 20 m, Lake Constance biomass trajectories were broadly similar to trajectories observed in the Great Lakes across comparable depth strata.

**Figure 5.**
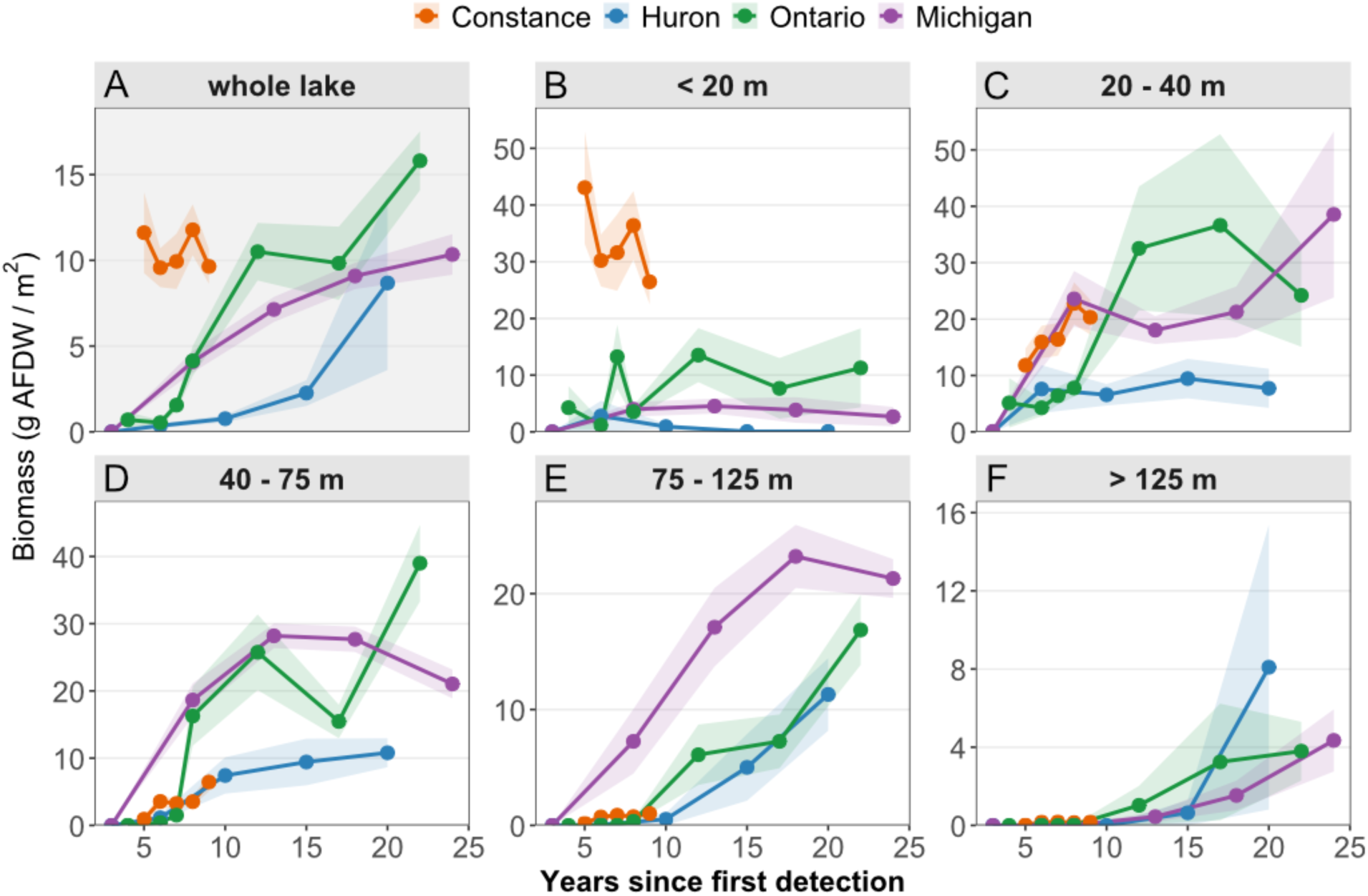
Mean biomass of quagga mussels > 5 mm shell length in Lake Constance and three Laurentian Great Lakes since first detection. Biomass is expressed as ash-free dry weight (AFDW). (A) Lake-wide means weighted by the proportion of three-dimensional lakebed area within each depth stratum. (B-F) Means by depth stratum for Lake Constance, Lake Huron, Lake Ontario, and Lake Michigan. Ribbons indicate standard errors.

## Discussion

Five years of lake-wide monitoring reveal rapid shallow-water saturation and ongoing deep-water expansion of quagga mussels in Lake Constance. The general sequence of rapid littoral colonization followed by slower expansion into deeper strata is consistent with patterns observed in the Laurentian Great Lakes (Karatayev and Burlakova 2025b). However, our expectation of lake-wide exponential increases in density and biomass, motivated by the projections of Kraemer et al. (2023), was not supported by the extended 2021-2025 time series. Instead, shallow-water populations appear to have reached carrying capacity before the first sampling year, while densities continue to increase at intermediate depths. This depth-specific pattern indicates that whole-lake means can obscure ongoing invasion dynamics in deeper habitats and that early projections based on short time series should be interpreted in light of depth-specific population states.

The dataset includes methodological changes in the first year of sampling. However, the comparison between BIS images and Ponar samples indicates that density estimates from the two methods were comparable for mussels > 5 mm shell length. Excluding mussels < 5 mm underestimates density, especially in deeper strata where small individuals are common, but has a smaller effect on biomass because very small mussels contribute little to total AFDW (Karatayev et al. 2021a). These limitations should be considered when interpreting absolute values, but they are unlikely to alter the main depth-specific trends. The study therefore provides a robust temporal record of quagga mussel invasion dynamics in a deep European lake and a foundation for continued monitoring.

### Shallow populations are stable

Shell-length distributions in the shallowest stratum suggest declining recruitment relative to growth of established individuals. In waters < 20 m, skewness declined sharply from 2023 to 2025, indicating a shift toward larger individuals and a reduced contribution of small mussels. Mann-Kendall tests showed no significant increase in shallow density or biomass, and station-level models showed no significant biomass increase at any station < 20 m. Because shallow regions contain the largest individuals and the highest biomass, they dominate lake-wide biomass estimates and can mask expansion occurring at depth.

### Expansion continues at intermediate depths

Significant monotonic increases in density between 40 and 125 m depth show that the invasion is still actively progressing beyond the littoral zone. The absence of significant trends below 125 m probably reflects the early stage and slow growth of deep-water populations rather than a lack of recruitment. Cold, food-limited conditions in the hypolimnion reduce growth rates, making changes harder to detect over a five-year interval (Karatayev and Burlakova 2025c). The predominance of positive station-level slopes at depths > 75 m supports the interpretation of gradual expansion toward profundal habitats. Based on similarities with the Great Lakes, biomass at these depths is likely to continue increasing over coming decades.

### European and North American trajectories differ

Lake Constance differs from the Great Lakes in the magnitude and timing of shallow-water biomass accumulation. Although quagga mussels arrived in the Great Lakes almost two decades earlier than in central Europe, shallow-water biomass in Lake Constance is already substantially higher, causing lake-wide biomass to exceed Great Lakes levels at comparable time after first detection. In the Great Lakes, quagga mussel biomass at shallow depths started to decline 13 to 15 years after first detection (Karatayev abd Burlakova, 2025b), whereas Lake Constance appears to have already reached shallow-water saturation six years after first detection. Delayed detection may partly explain this apparent rapidity: the first reported quagga mussels in Lake Constance were already large, suggesting that the species most likely established before 2016.

A true difference in invasion trajectory is also plausible. Lake area and morphometry strongly influence *Dreissena* invasion dynamics, and the much larger surface area and fetch of the Great Lakes likely increase wave disturbance in shallow habitats relative to perialpine lakes (Karatayev et al. 2021b; Karatayev et al. unpubl.). Therefore, it is possible that weaker shallow-water disturbance in Lake Constance may allow higher littoral biomass. At the same time, ongoing re-oligotrophication and reduced full-depth mixing complicate attribution of pelagic ecosystem change to mussel filtration alone (Fink et al. 2015; Murphy et al. 2018). These interacting drivers limit direct transfer of Great Lakes predictions to European deep lakes.

The recent 18-lake synthesis provides a useful context for this contrast: *Dreissena* effects on Secchi depth, chlorophyll, phytoplankton, zooplankton, and phosphorus generally appeared quickly in inland lakes, were delayed but stronger in the Laurentian Great Lakes, and were unusually weak or difficult to detect in Lakes Constance and Geneva despite substantial quagga mussel biomass (Karatayev et al. unpubl.). This broader pattern supports our interpretation that Lake Constance is not simply a smaller analogue of the Great Lakes, but a perialpine system in which high shallow-water mussel biomass, ongoing deep-water expansion, re-oligotrophication, and changing mixing interact.

### Deep expansion and lake-wide consequences

Future whole-lake quagga mussel biomass in Lake Constance remains uncertain. If shallow-water biomass has overshot carrying capacity, future declines in the littoral zone could offset biomass increases at depth. Conversely, if perialpine lakes can sustain higher biomass than the Great Lakes across depth strata, whole-lake biomass could increase substantially as colonization progresses. Because more than 60% of the lakebed area lies below 45 m, even moderate biomass increases in deeper strata could strongly affect whole-lake mussel biomass.

The ecological implications of increasing deep-water biomass are potentially large. As quagga mussels expand into profundal habitats, filtration pressure and biodeposition may increasingly influence nutrient cycling, benthic-pelagic coupling, and the deep-water food web. In the Great Lakes, deep-water colonization has been associated with reduced phytoplankton biomass, altered zooplankton communities, and cascading effects on pelagic fish (Higgins and Vander Zanden 2010; Mayer et al. 2014; Karatayev and Burlakova 2025c). Whether effects of similar magnitude will emerge in perialpine European lakes remains uncertain; the current cross-lake evidence suggests that pelagic responses in Lakes Constance and Geneva may be masked or modified by climate-driven mixing changes and re-oligotrophication (Karatayev et al. unpubl.). Continued standardized monitoring of mussel populations, nutrients, phytoplankton, and zooplankton will be essential to separate these drivers and improve projections (Flämig et al. 2024).

### Implications for European lakes

Lake-specific features such as productivity, stratification, mixing regime, substrate, and morphometry shape quagga mussel invasion trajectories (Karatayev et al. 2021b; Karatayev and Burlakova 2025c). Consequently, observations from Lake Constance should not be transferred to other perialpine lakes without careful consideration. Longer time series across multiple lakes will be needed for robust forecasting, especially because the present dataset captures only an early segment of a long invasion process. Nevertheless, Lake Constance trajectories show that deep European lakes can experience rapid shallow-water saturation while deeper habitats remain in active expansion. This pattern should be incorporated into monitoring programs, ecosystem models, and management planning for European lakes increasingly exposed to quagga mussel invasion.

## Acknowledgments

This study was financed by the grants “SeeWandel: Life in Lake Constance - the past, present and future” and “SeeWandel-Climate: Modelling the consequences of climate change and neobiota for Lake Constance” within the framework of the Interreg V and Interreg VI programmes “Alpenrhein-Bodensee-Hochrhein (Germany/Austria/Switzerland/Liechtenstein)” which funds are provided by the European Regional Development Fund as well as the Swiss cantons (SeeWandel, SeeWandel-Climate), and the Swiss Confederation (SeeWandel). Additional financial support was provided by the International Commission of Lake Constance Water Conservation (Internationale Gewässerschutzkommission für den Bodensee, IGKB). The funders had no role in study design, data collection and analysis, the decision to publish, or preparation of the manuscript. Data collection was made possible by the boat crews of the Institute for Lake Research (ISF) of the State Institute for the Environment Baden-Württemberg (LUBW) and the University of Konstanz. We thank Silvan Rossbacher for coordinating and conducting the first two monitoring campaigns, and Raphael Bossart, Silvana Käser, Lale Baehni, Christoph Walcher, Antonia Zuber, Rebecca Dorendorf and Josephine Alexander for assistance during field work. We are grateful to Lydia Seitz and Nadine Jabornegg for their help with data preparation and analyses. Generative artificial intelligence (Claude Sonnet 4.6 (Anthropic) and ChatGPT - 5.5 (OpenAI)) were used to assist with writing and tidying of R scripts, and for outlining and grammar correction during manuscript drafting. All content was reviewed by the authors, who take responsibility for the final manuscript. No conflicts of interest are declared.

## Data availability statement

The data, metadata, and analysis code supporting the results have been archived on Zenodo and are available at: https://doi.org/10.5281/zenodo.20763368.

