## Supplementary Figures and Tables for "Rapid shallow-water saturation and deep-water expansion of an invasive freshwater ecosystem engineer in a deep European lake"

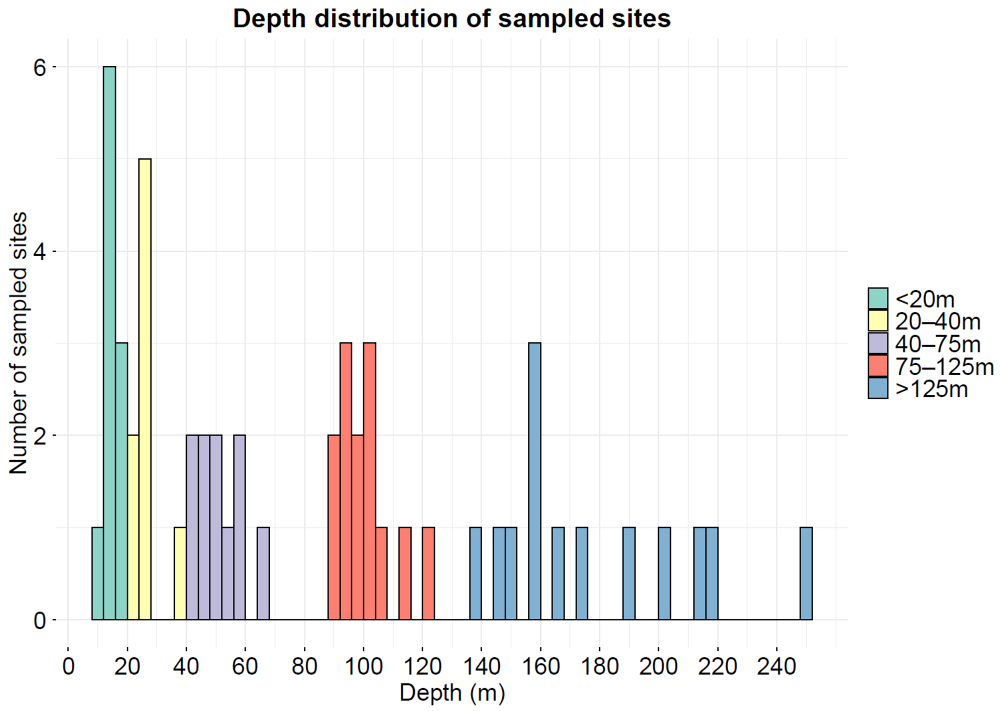


**Figure S1. Depth distribution of all sampling stations.** Colors indicate the five depth strata used in the analyses.

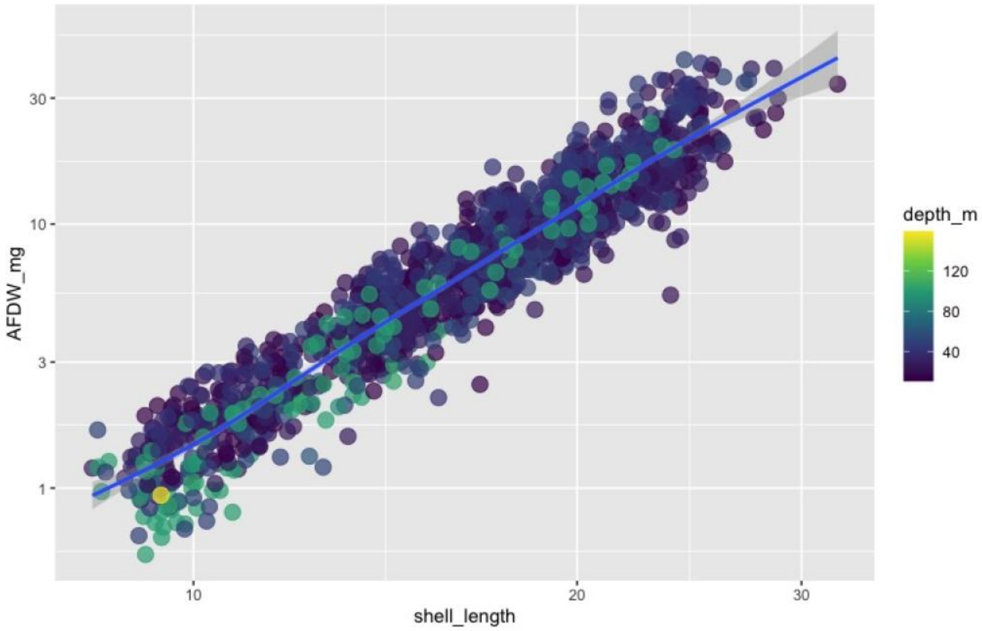


**Figure S2. Relationship between measured shell length and ash-free dry weight (AFDW) for 761 quagga mussels sampled in Lake Constance in 2022.** Each point represents one individual, and color indicates depth of origin. This regression was used to estimate the coefficients in Table S2 and Formula S1. Both axes are nonlinear.

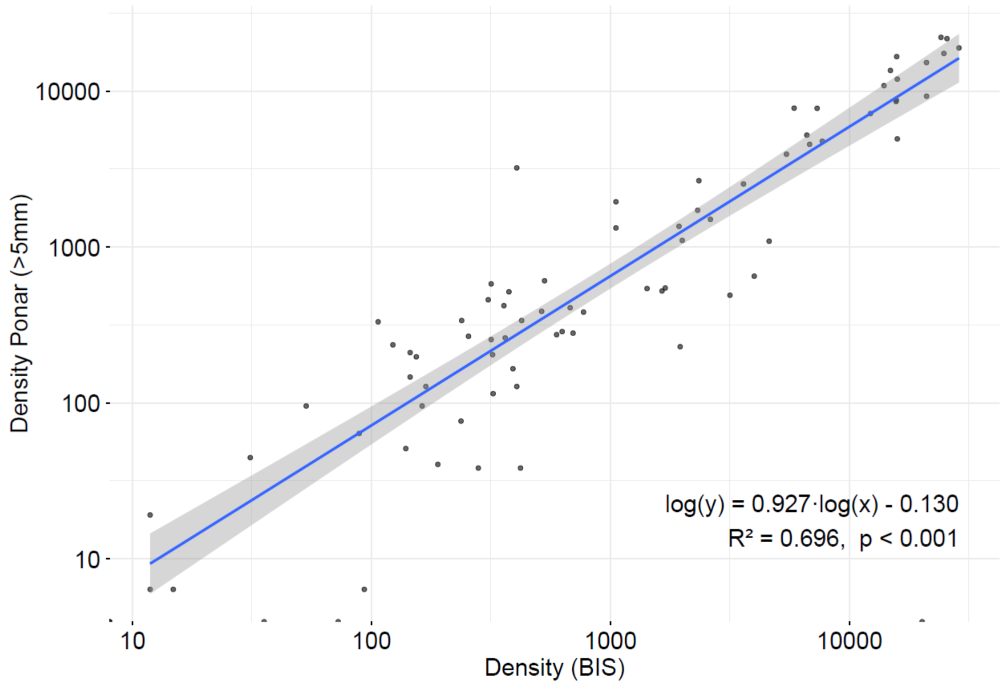


**Figure S3. Relationship between quagga mussel densities estimated from Ponar samples and the Benthic Imaging System (BIS) in 2021 and 2022.** Each point represents the mean of three subsamples at one station (n = 80). Ponar density includes only mussels > 5 mm shell length.

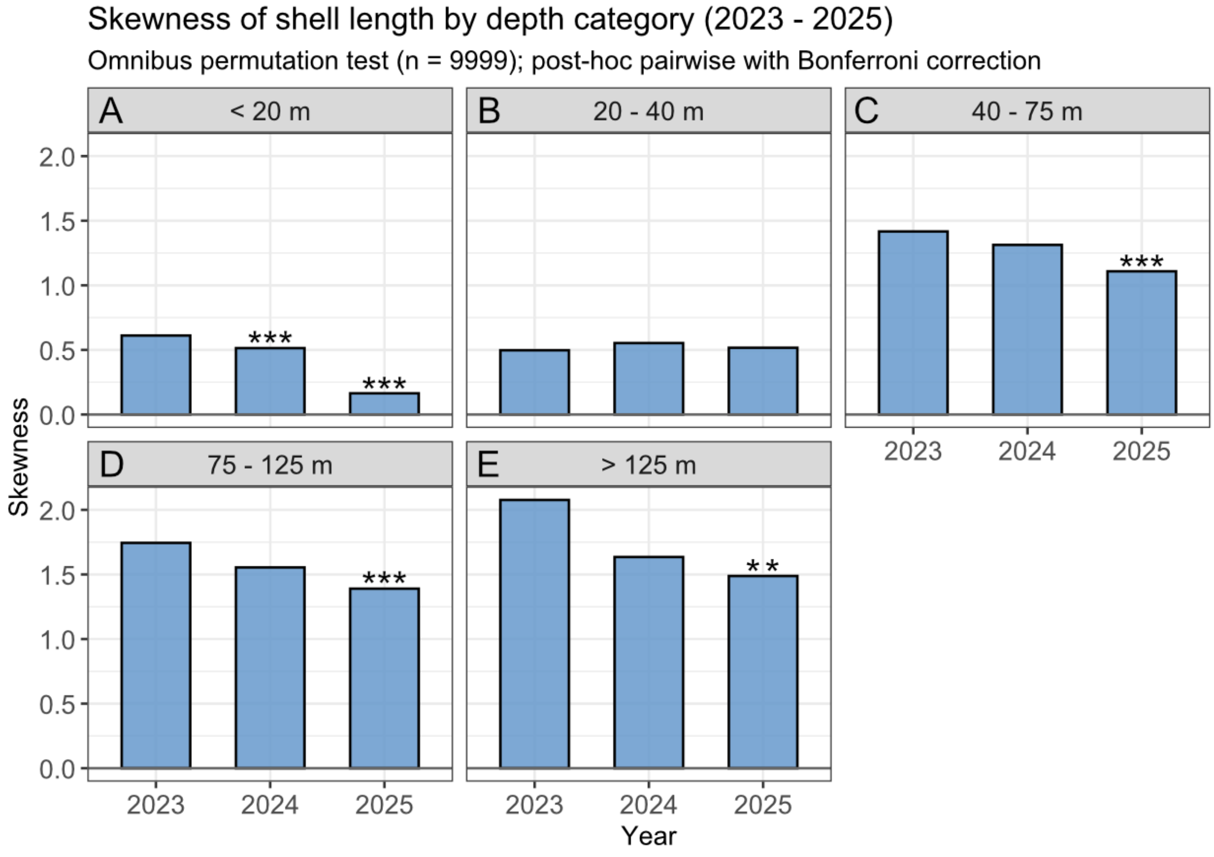


**Figure S4. Skewness of shell-length distributions in 2023, 2024, and 2025 by depth stratum.**

Stars indicate significant differences relative to 2023 after Bonferroni correction (** $p < 0.01$; *** $p < 0.001$).

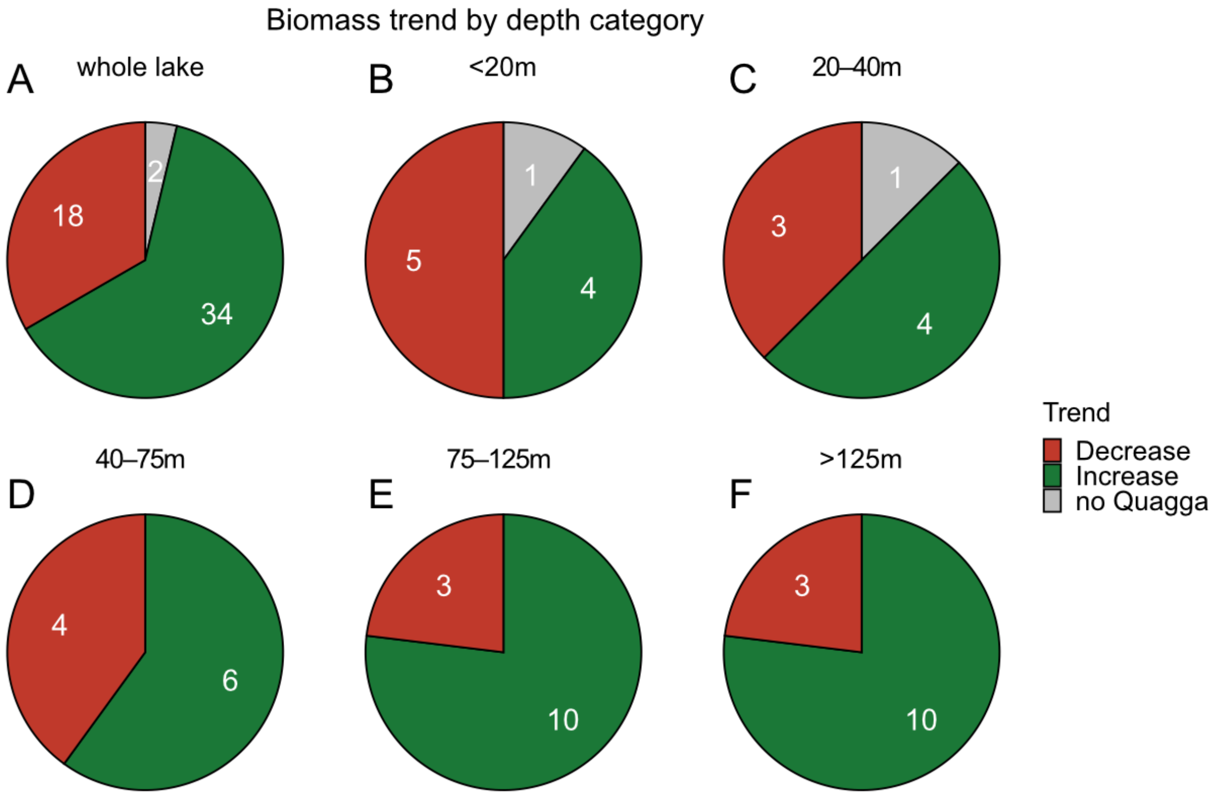


**Figure S5. Proportion of stations with positive and negative biomass slopes over the sampling period, regardless of statistical significance.** Panels show trends for the whole lake (A) and for each depth stratum (B-F). The two stations that remained quagga-free are shown in gray.

**Table S1.** Coordinates and target depth for each station and years sampled with Ponar grabs.

| station | latitude | longitude | depth | depth category | years monitored |
|---|---|---|---|---|---|
| BQ | 47.52019 | 9.72192 | 60 | 40-75m | 2021-2025 |
| EA | 47.6093 | 9.515232 | 12.4 | <20m | 2021-2025 |
| EB | 47.60578 | 9.511395 | 40.5 | 40-75m | 2022-2025 |
| EC | 47.60219 | 9.505534 | 56.3 | 40-75m | 2022-2025 |
| ED | 47.59795 | 9.498636 | 104.6 | 75-125m | 2021-2025 |
| EE | 47.563 | 9.448114 | 191.9 | >125m | 2021-2025 |
| EF | 47.55158 | 9.431291 | 91.2 | 75-125m | 2022-2025 |
| EG | 47.54574 | 9.42266 | 64.5 | 40-75m | 2021-2025 |
| EH | 47.5412 | 9.416509 | 37.9 | 20-40m | 2021-2025 |
| EI | 47.53721 | 9.41136 | 9.8 | <20m | 2022-2025 |
| EQ | 47.58535 | 9.477579 | 219.9 | >125m | 2022-2025 |
| FA | 47.66124 | 9.404329 | 19.7 | <20m | 2022-2025 |
| FB | 47.66043 | 9.40363 | 23.7 | 20-40m | 2021-2025 |
| FC | 47.65536 | 9.40045 | 48.1 | 40-75m | 2022-2025 |
| FD | 47.64782 | 9.39359 | 101.7 | 75-125m | 2021-2025 |
| FE | 47.63568 | 9.383905 | 203 | >125m | 2022-2025 |
| FF | 47.61762 | 9.369486 | 156.4 | >125m | 2022-2025 |
| FG | 47.59991 | 9.35611 | 100.6 | 75-125m | 2021-2025 |
| FH | 47.59348 | 9.350648 | 46.8 | 40-75m | 2022-2025 |
| FI | 47.58971 | 9.347552 | 24.9 | 20-40m | 2022-2025 |
| FJ | 47.58857 | 9.34656 | 14.2 | <20m | 2021-2025 |
| FQ | 47.62396 | 9.375571 | 250.1 | >125m | 2021-2025 |
| LA | 47.52451 | 9.626492 | 49.7 | 40-75m | 2022-2025 |
| LB | 47.51992 | 9.583918 | 92.4 | 75-125m | 2022-2025 |
| LC | 47.50041 | 9.593647 | 24.1 | 20-40m | 2022-2025 |
| LQ | 47.53496 | 9.641691 | 55.7 | 40-75m | 2021-2025 |
| MA | 47.68932 | 9.264854 | 156.7 | >125m | 2021-2025 |
| MB | 47.66692 | 9.237725 | 100.8 | 75-125m | 2022-2025 |
| MC | 47.6601 | 9.241741 | 92.6 | 75-125m | 2021-2025 |
| MD | 47.65656 | 9.243229 | 92.6 | 75-125m | 2022-2025 |
| ME | 47.65061 | 9.218483 | 47.7 | 40-75m | 2022-2025 |
| MF | 47.64682 | 9.212718 | 25.6 | 20-40m | 2021-2025 |
| MQ | 47.68237 | 9.256255 | 175.4 | >125m | 2021-2025 |
| QA | 47.81561 | 9.041744 | 25.4 | 20-40m | 2021-2025 |
| QB | 47.79645 | 9.070255 | 90.5 | 75-125m | 2021-2025 |
| QC | 47.78319 | 9.094511 | 112.3 | 75-125m | 2022-2025 |
| QD | 47.75945 | 9.13444 | 146 | >125m | 2022-2025 |
| QE | 47.71218 | 9.212517 | 123.2 | 75-125m | 2022-2025 |
| QF | 47.65812 | 9.291947 | 212.6 | >125m | 2022-2025 |
| UA | 47.678 | 9.100478 | 12.5 | <20m | 2021-2025 |
| UB | 47.6751 | 8.99694 | 41.4 | 40-75m | 2022-2025 |
| UC | 47.64836 | 8.907392 | 26.4 | 20-40m | 2021-2025 |
| UD | 47.64636 | 8.909202 | 14.2 | <20m | 2022-2025 |
| UE | 47.72395 | 8.973944 | 22.4 | 20-40m | 2021-2025 |
| UF | 47.72913 | 8.981217 | 14.3 | <20m | 2022-2025 |
| UG | 47.70766 | 9.063137 | 19.8 | <20m | 2022-2025 |
| WA | 47.56435 | 9.623937 | 14.6 | <20m | 2021-2025 |
| WB | 47.56106 | 9.613892 | 97.1 | 75-125m | 2022-2025 |
| WC | 47.54595 | 9.568071 | 148.2 | >125m | 2022-2025 |
| WD | 47.518 | 9.485308 | 164.6 | >125m | 2021-2025 |
| WE | 47.51696 | 9.479989 | 159.6 | >125m | 2021-2025 |
| WF | 47.51482 | 9.474363 | 98.8 | 75-125m | 2021-2025 |
| WG | 47.5103 | 9.451038 | 19.4 | <20m | 2021-2025 |
| WQ | 47.55628 | 9.599434 | 139.7 | >125m | 2022-2025 |

**Table S2.** Shell-length-specific coefficients for ash-free dry weight (AFDW) calculation from mussel size, used in Formula S1. Coefficients were estimated from the regression shown in Fig. S2.

| Depth category | a | b |
|---|---|---|
| < 30 m | - 6.00895 | 2.81329 |
| 0 - 50 m | - 6.34713 | 2.956302 |
| 50 - 90 m | - 6.92735 | 3.15219 |
| > 90 m | - 7.45429 | 3.313763 |

**Table S3.** R packages used in this study, grouped by purpose.

| Purpose | R packages |
|---|---|
| Data manipulation | broom; data.table; tidyverse |
| Plot generation | ggh4x; ggrepel; ggridges; tidyverse |
| Map generation and bathymetric data processing | rnaturalearth; terra |
| Statistical testing | moments; trend |

**Formula S1.** Relationship between quagga mussel ash-free dry weight (AFDW) and shell length. AFDW is in mg, shell length is in mm, and depth-specific coefficients a and b are listed in Table S2.

$$\log(\text{AFDW}) = a + b \times \log(\text{shell length})$$
